# Biodecomposing *Spirulina platensis* by a *de novo* designed *Bacillus subtilis*-based method to develop a medium for the high cell-density cultivation of *Escherichia coli* in batch mode

**DOI:** 10.1101/2023.06.30.547221

**Authors:** Xiaowen Li, Zhengqi Wei, Jingyi Ge, Yingying Pan, Xiang Chu, Baolin Huang, Jiawei Zhao, Yanzhao Li, Yizhuang Zhou

## Abstract

High cell-density cultivation (HCDC) is fundamental to basic research and industrial applications, especially in batch mode. However, limited media are available for batch culture of HCDC, because the media for batch culture must contain extremely sufficient nutrients on the one hand and few or even no substrates to generate detrimental metabolite on the other hand to attain HCDC. *Spirulina platensis* (SP), a new media material, is considered ideal for the development of such media. Here, we develop a biolysis method for SP degradation based on the cultivation supernatant of *Bacillus subtilis* and extensively demonstrate its higher degradation (indicated by the production of more small peptides and free amino acids) and cultivation effectiveness with three other methods. Based on its SP hydrolysates, a modified SP-based broth (MSP) is then formulated. Tests on *Escherichia coli* K-12 show that MSP achieves HCDC with several benefits: (*i*) its maximum optical density at 600 nm is ∼16.67, significantly higher than that of Luria-Bertani (LB) broth (only ∼6.30); (*ii*) MSP requires only 36 h to reach peak growth, much faster than that of LB (48 h); (*iii*) its maximum growth (1.12 ± 0.01 h^−1^) is significantly higher than that of LB (0.20 ± 0.00 h^−1^); (*v*) MSP initiates growth immediately after inoculation (lag time <0), comparable to LB; (*iv*) the number of viable cells in MSP is high (∼2.16 × 10^11^ ml^-1^), ∼10.19 times the amount in LB. Consequently, we envision MSP will become the first choice for *E. coli* HCDC batch culture in the future.

**Importance:** So far, it is the first time to develop a high-efficiency method for transforming *Spirulina platensis* (SP) into medium ingredients. Based on its SP hydrolysates, a high cell-density cultivation (HCDC) medium for the batch culture of *Escherichia coli* is formulated for the first time, which is greatly beneficial for both basic research and industrial applications. In addition to HCDC, the SP hydrolysates can be extended to a wide range of applications, due to their rich nutrient content. Besides, this study demonstrates for the first time that SP is an ideal material to develop HCDC media. Furthermore, this study demonstrates that medium development and modification for batch culture can attain HCDC, without the development of new culture technologies. Therefore, this study highlights the importance of the rebirth of medium development and modification and supports the shift from developing new culture technologies to medium development and modification for HCDC in batch mode.

## Introduction

Cell growth is fundamental to both basic research and industrial application, requiring bacteria to grow to high cell density to maximize biomolecules including plasmids and proteins, especially for the pioneer bacterium *Escherichia coli*. To achieve high cell-density cultivation (HCDC), requisite nutrients should be sufficiently supplied, and toxic or inhibitory factors generated by metabolism amid the cultivation should be avoided or removed meanwhile (1). Under this logic, some promising culture technologies, including fed-batch culture (2, 3), dialysis culture (4-6), and semicontinuous culture (7), had been developed and applied accordingly, attaining HCDC *via* providing additional nutrients and diluting or removing inhibitors coincidentally. However, they all require sophisticated instrumentation and some of them such as dialysis culture waste a considerable amount of nutrients, combinedly leading to high cost. Therefore, they are not routinely used in laboratories and industries. In contrast, batch culture is preferred in laboratories and even in industry, owing to simple handling, high feasibility, easy parallelization, and reduced investment in equipment. To attain HCDC for batch culture, the media should contain extremely sufficient nutrients on one hand, as all necessary nutrients are provided once before culture in batch mode; on the other hand, the media should contain few or even no substrates to generate detrimental metabolites, due to no removal or dilution of detrimental metabolites during the whole cultivation. Accordingly, it is difficult for a medium to satisfy both requirements together. Therefore, to the best of our knowledge, such a medium is not yet available. For example, the widely-used rich Luria-Bertani (LB) medium can only reach saturation of ∼7 optical density at 600 nm (OD600) for *E. coli* (8); glucose, one most favorable carbon source for *E. coli* (9), inhibits growth above the concentration of 50 g/l (10) or by its metabolite acetate acid (11). In this sense, developing a novel medium for HCDC is of pronounced importance and urgent.

*Spirulina platensis* (SP), a member of the Phormidiaceae family with rich nutrients (12), is an ideal material to potentially develop culture media for HCDC based on the following considerations. First, it is one of the richest protein sources with a protein content of around 60-70% (13), greatly higher than that in egg, beer yeast, skimmed powdered milk, fish, soy, and beef meat (14), some of which are traditional sources for media. Second, the most abundant monosaccharide in SP is rhamnose, which accounts for 53% of the total sugars (15), while glucose is relatively low (<2%) (16); this is beneficial to prevent/decrease acetate acid generation, especially by excess glucose (17). Third, SP contains all essential minerals and trace elements (14, 18), as well as luxuriant bioavailable vitamins including vitamins A, E, B1, B7, and B8 (14). Finally, SP contains numerous antioxidant ingredients, including phycocyanin, carotenoids, and xanthophylls (19), which may be beneficial to ameliorate oxidant stress under aerobic conditions. Besides, SP, as a photoautotrophic microorganism, can be cultivated on a large scale in a small area at low cost and absorbs atmospheric CO_2_, making a positive impact on the environment (20). Therefore, SP was here selected to develop an HCDC medium for *E. coli*.

So far as we know, little or almost no research has been reported on transforming SP into medium ingredients (20, 21), hampering its utilization for HCDC or even routine cultivation. The difficulty of its transformation may suffer from the two following reasons. First, SP cell is difficult to disrupt due to their high resistance, especially for dry SP (22). Second, SP proteins should be further degraded into small peptides or even free amino acids for HCDC, as most bacteria can only utilize free amino acids due to the scarcity of peptide transporters (23). Even for *E. coli* with several oligopeptide permeases and peptidases (8), efficient degradation is also beneficial, as free amino acids can be directly utilized to accelerate growth, with small peptides instead of large peptides utilized after complete degradation. *Bacillus subtilis* is commonly found in the soil. Provided that nutrients are sparse or changing in soil, *B. subtilis* has to produce extracellular proteases to support its growth and survive *via* protein degradation. So far, at least eight secreted proteases have been reported (24). Except for the cell wall-associated protease (WprA) (25), the remaining proteases are secreted into the growth medium, potentially supplying an opportunity to efficiently degrade SP by using the cultivation supernatant of *B. subtilis*.

Here, we developed a *B. subtilis*-based method to lyse SP and demonstrated that this method outperforms the three mechanical methods in terms of small peptides obtained, amino acids liberated, and growth performance achieved. Based on the products of this method, as a test for *E. coli* K-12, we further developed a modified SP-based medium (hereinafter named MSP) and found that MSP achieved HCDC (OD600 of ∼16.97) rapidly within 36 h, with a significantly larger maximum growth rate and ∼10.19 times the number of viable cells in comparison to the reference LB medium. To the best of our knowledge, it is the first time for us to biodegrade SP to achieve HCDC in batch mode. We envisage that MSP developed here will become the medium of first choice for *E. coli* HCDC in batch culture in the future.

## Results

### Development of biolysis methods using varying cultivation supernatant of *B. subtilis*

As all the 7 aforementioned extracellular proteases secreted into cultivation supernatant are highly produced during the plateau growth phase (24), the cultivation supernatant sampled at 72 h was used for SP degradation, based on the growth kinetics of *B. subtilis* determined here (Fig. S1). It was observed that *B. subtilis* growth was significantly enhanced in Nutrient Broth (NB) than in LB (Fig. S1), which may provide more extracellular proteases in NB than in LB to increase SP degradation. However, both NB and LB cultivation supernatants were explored to see which one was indeed better. To further determine the optimal volume, aliquots of 1 ml, 3 ml, and 9 ml of cultivation supernatants were added into 300 ml sterilized SP solutions, each of which contained 45 g SP dry power with initial pH at ∼7 and was autoclaved at 121℃ for 15 min to eliminate the effect of microorganisms on SP consumption. After 5 h degradation, we found that the pH levels of all SP solutions were decreased (Fig. S2A), possibly due to the two highest-abundance acidic amino acids liberated into the solutions, namely glutamic acid (∼8.386%) and aspartic acid (∼5.793%) (14), which were confirmed by the results below (Fig. 4a). Among them, the pH fall was the largest in the product of the method with 3 ml NB cultivation supernatant (Fig. S2A), with a significant difference with the products of others except by using 9 ml NB cultivation supernatant, indicating that 3 ml NB cultivation supernatant may have the best lysis. To confirm it, a direct formol titration method (26) was applied. Expectedly, our results showed that 3 ml NB cultivation supernatant has the significantly best degradation effectiveness (Fig. S3A).

Considering that proteases may partially reduce or even lose their activities under acidic conditions, we wondered whether the addition of fresh cultivation supernatant could increase SP degradation. To this end, 4 extra successive addition-degradation cycles were performed. Due to pH falls, SP products were neutralized with 1 M NaOH to eliminate their effects on degradation, and then added with 1.893 g of KH_2_PO_4_ and 2.803 g of K_2_HPO_4_ (0.1 M potassium phosphate, pH 7.0) to provide buffering capacity to postpone pH falls. Subsequently, fresh cultivation supernatant with respective volume was added and then subjected to 5-h degradation. It was observed that the pH levels of all SP products were always decreased after each 5-h degradation and the processing with 3 ml NB cultivation supernatant regularly yielded the largest pH falls, albeit with or without significant difference compared to other processing (Figs. S2B-2E). Subsequent formol titration assays repeatedly demonstrated that 3 ml NB cultivation supernatant significantly outperformed all other treatments (Figs. S3B-3D). After 25-h degradation, the processing with 3 ml NB cultivation supernatant achieved the best degradation performance (Fig. 1A).

**Fig. 1.**
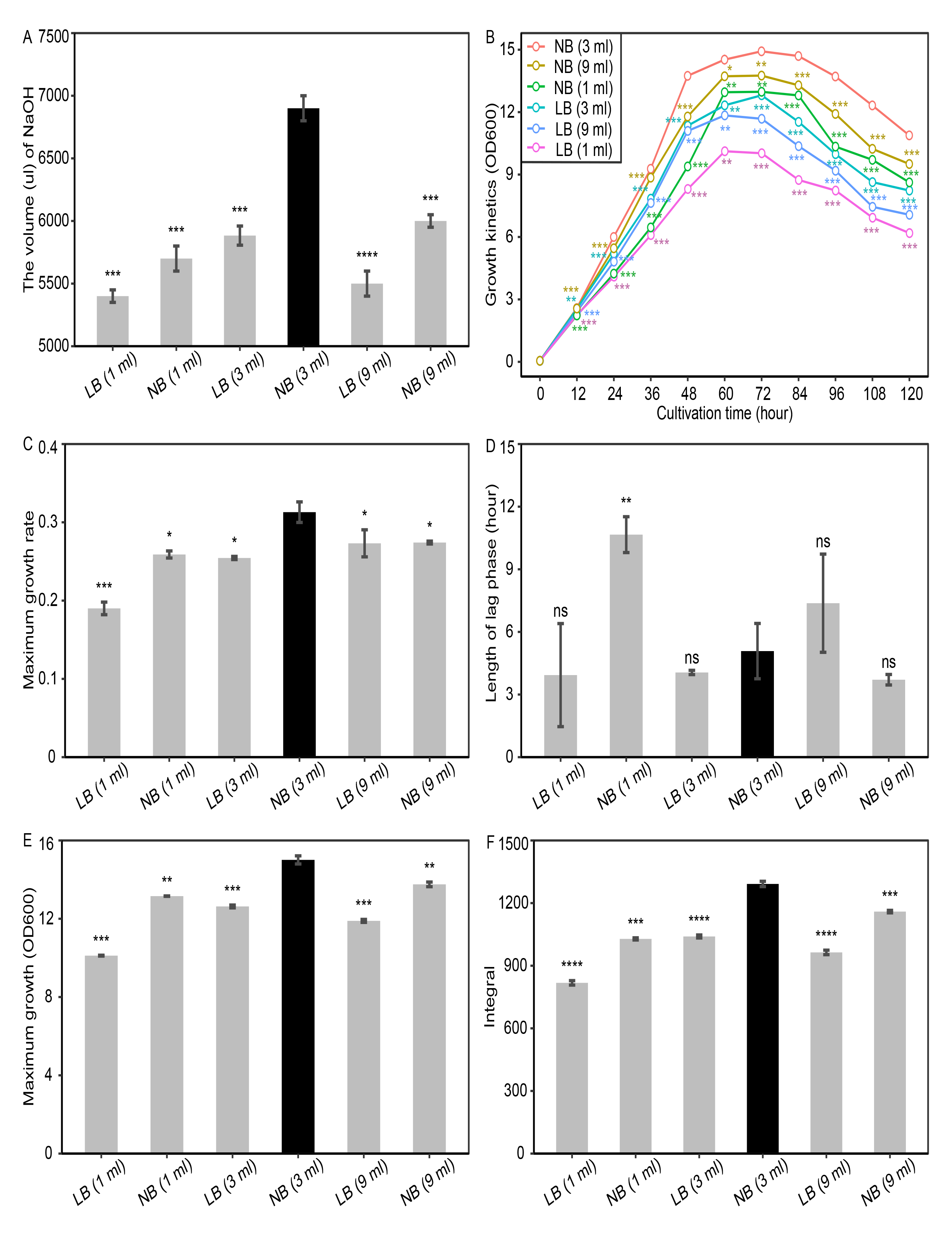
Developing a biolysis method based on *B. subtilis* cultivation supernatant. Varying volumes of *B. subtilis* supernatant cultivated in NB and LB were compared to develop the biolysis method. (A), the final overall free amino acids after 25-h degradation measured by formol titration. The volume (ul) of 0.1 M NaOH required to reach the endpoint titration indicates biodegradation efficiency. The growth kinetics of *E. coli* K-12 in the resultant SP broths (B), and 4 growth parameters including maximum growth rate (C), length of lag phase (D), maximum growth (E), and integral (F). \**P* <0.05, \*\**P* <0.01, \*\*\**P* <0.001, \*\*\*\**P* <0.0001, and ns, *P* >0.05, t-test compared with the SP product generated by using 3 ml NB cultivation supernatant; value and error bar represent mean ± standard deviation from biological triplicate experiments.

### Comparison of growth performance on original SP broths as culture media

After the above 25-h biodegradation, we accordingly obtained six different original SP broths. To further determine which one was best, a quantitative comparison of cultivation was conducted *via* testing *E. coli* K-12. It was observed that the best growth was significantly achieved for the broth derived from the degradation by using 3-ml NB cultivation supernatant regardless of the sampled times (Fig. 1B), with a significantly highest maximum growth rate (Fig. 1C) and a significantly or insignificantly reduced lag time (Fig. 1D). Also, this broth attained a significantly highest maximum OD600 predicted by *grofit* (Fig. 1E) and obtained a statistically highest integral value on the whole (Fig. 1F). Consequently, based on all findings (Fig. 1), the above processing with 3 ml NB cultivation supernatant was determined as the protocol of the biolysis method developed in this study and the resultant original SP broth (OSP) was obtained for following experiments.

### Superiority in degradation and cultivation effectiveness compared with other methods

Although there are several conventional methods available to disrupt SP cells for the extract of phycocyanin (27), little or almost no methods are available to degrade SP for developing the SP-based media. It is worth mentioning that the methods used for the extract of phycocyanin are relatively different from the SP degradation in this study, as the former involves gentle cell disintegration to obtain intact phycocyanin, while the latter involves vigorous SP degradation into short peptides and even free amino acids. However, in an attempt to demonstrate the conspicuousness of our method, we compared the biolysis method with three conventional methods including alternate freezing and thawing (28), sonication (29), and water bath.

For comparison of degradation effectiveness, formol titration assays were carried out. Our findings showed that the released amino acids are greatly significantly higher in the products of our biolysis method than in the products of the three conventional methods (*P* < 0.0001, Fig. 2A), demonstrating that our method is more effective. To further verify its superiority, cultivation trials on *E. coli* K-12 were performed and monitored from 12 h to 120 h. It was observed that the cell densities reached in OSP generated by our biolysis method were significantly higher than those in other SP broths during the whole incubation period, with an averaged maximum OD600 of ∼14.92 at 72 h, while only approaching 10.36, 6.90, and 6.90 for alternate freezing and thawing, sonication and water bath at 60 h respectively (Fig. 2B). *E. coli* K-12 was cultivated with a maximum growth rate of 0.3130 ± 0.0131 h^-1^, significantly higher (*P* < 0.001) than that in the SP broths generated by the alternate freezing and thawing (0.2355 ± 0.0017 h^-1^), sonication (0.1772 ± 0.0012 h^-1^) and water bath (0.1780 ± 0.0017 h^-1^) respectively (Fig. 2C). However, its lag time was significantly longer than those by sonication and water bath, but insignificantly shorter than that by alternate freezing and thawing (Fig. 2D), possibly illustrating why the time to reach maximum growth is longer than those by other methods (Fig. 2B). The maximum OD600 predicted by *grofit* was 15.0072 ± 0.2075, 10.1892 ± 0.0686, 8.1229 ± 0.1071, and 8.0278 ± 0.0780 for SP broths generated by the methods of biolysis, alternate freezing and thawing, sonication and water bath respectively (Fig. 2E). Finally, our biolysis method yielded a significant higher integral value than others (Fig. 2F). Taken together, these findings demonstrated that our biolysis method outperforms other methods mentioned here to achieve HCDC by testing *E. coli* K-12, albeit with a significantly or insignificantly longer lag time.

**Fig. 2.**
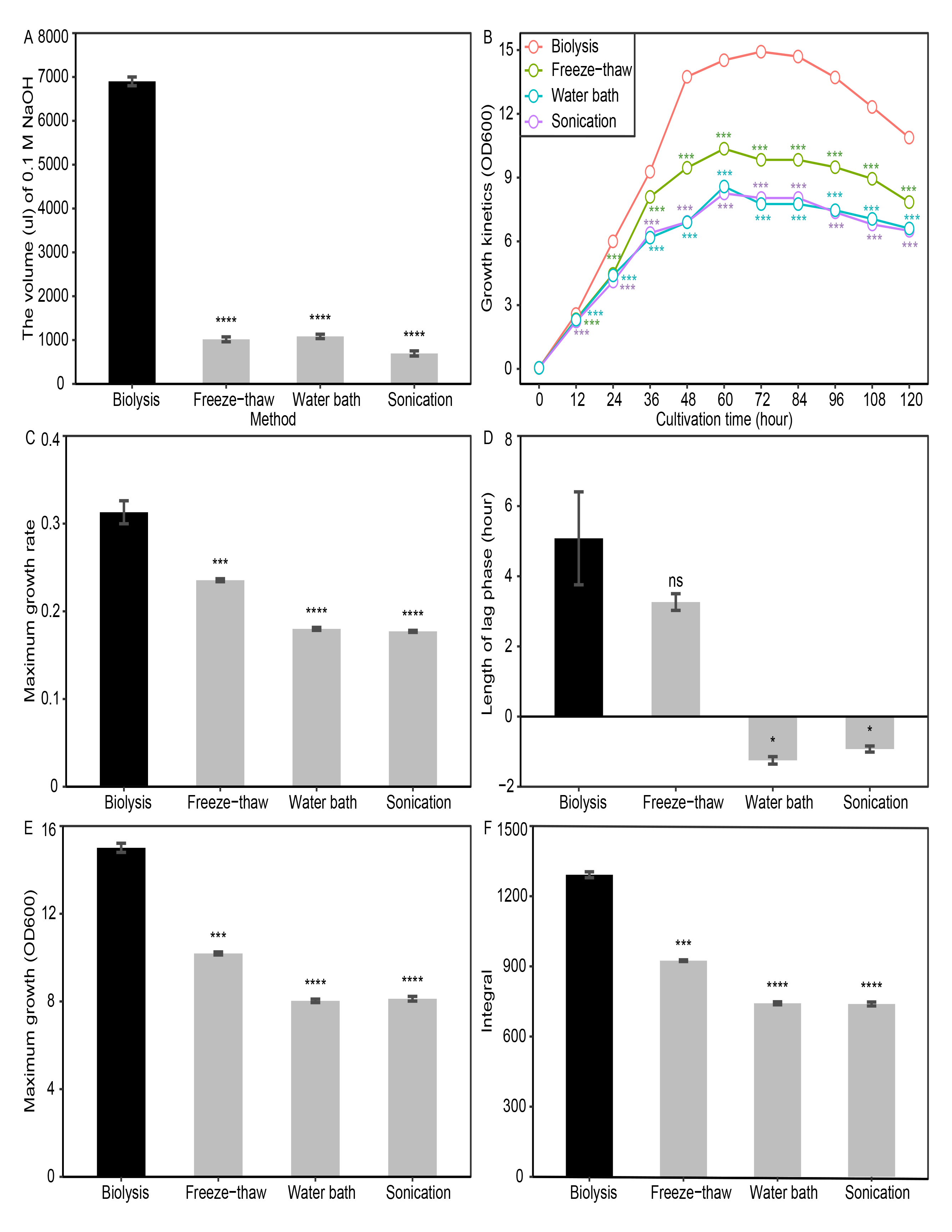
Comparison of 4 degradation methods in terms of degradation efficiency and growth performance. (A), the degradation efficiency measured by formol titration. The volume (ul) of 0.1 M NaOH required to reach the endpoint titration indicates degradation efficiency. For details, please refer to Materials and methods. The growth kinetics of *E. coli* K-12 in the SP broths generated by the 4 methods (B), as well as 4 extracted growth parameters including maximum growth rate (C), length of lag phase (D), maximum growth (E), and integral (F). \**P* <0.05, \*\*\**P* <0.001, \*\*\*\**P* <0.0001, and ns, *P* >0.05, t-test compared with the biolysis method; all experiments were performed in triplicate; error bar, mean ± SD.

### Demonstration of increased degradation effectiveness by gel electrophoresis

If SP is effectively decomposed and then efficiently degraded, SP proteins should be greatly released and transformed into short peptides or even free amino acids afterward. Accordingly, Sodium dodecyl sulfate-polyacrylamide gel electrophoresis (SDS-PAGE) was conducted to compare the degradation effectiveness in terms of short peptides between the 4 methods. As shown in Fig. 3A, protein degradation was evidenced by gel smearing in the SDS-PAGE portrait. Our biolysis method seemed to display increased degradation effectiveness in relation to other methods, as visualized by the more degraded proteins accumulated at the bottom of the gel (Fig. 3A). To further quantify the degradation, densitometry by ImageJ was conducted. It was observed that the area intensity for <10 kDa accounted for ∼42.92% of the total intensities for the biolysis method, which was significantly higher than that for other methods (Fig. 3B). Also, the area intensities for the ranges of 10-15 kDa (Fig. 3C) and 15-20 kDa (Fig. 3D) were conspicuously statistically higher for the biolysis method than for other methods respectively. Conversely, the intensity for the remaining areas was greatly lower than that by other methods with a statistical difference (*P* < 0.01, Fig. 3E). In conclusion, these findings authenticated that the biolysis method achieved higher degradation effectiveness with respect to other methods. This is very beneficial for some bacteria such as *Lactobacillus amylovorus* (30), which prefers peptide-bound amino acids to free amino acids, and *E. coli* with several oligopeptide permeases and peptidases (8).

**Fig. 3.**
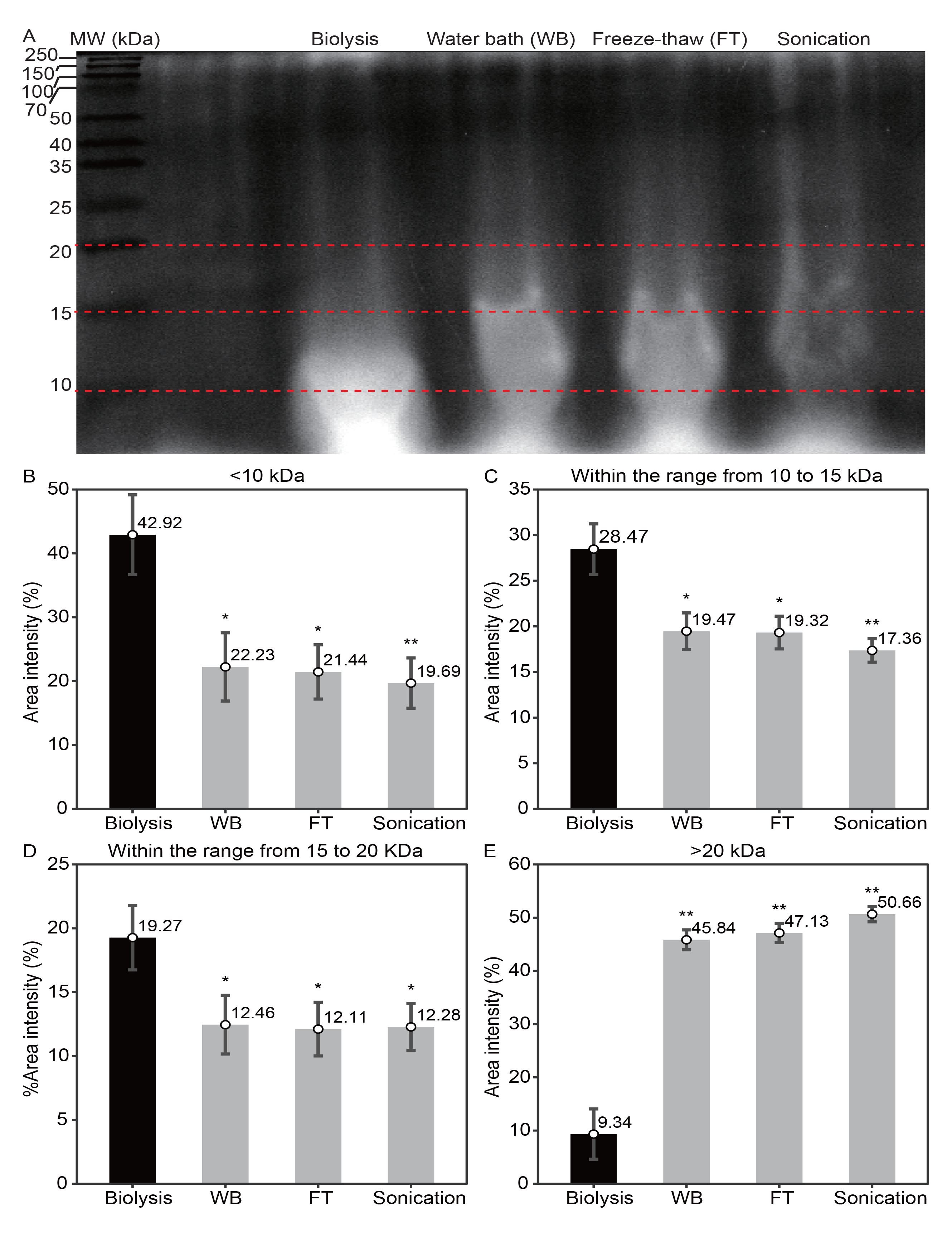
Demonstration of the elevated degradation effectiveness by our biolysis method in comparison to three mechanical methods by SDS-PAGE. SDS-PAGE profile of protein degradation by the 4 methods (A), as well as the percentage of area intensity for <10 kDa (B), the ranges of 10-15 kDa (C) and 15-20 kDa (D), and >20 kDa (E) respectively. \**P* <0.05, \*\**P* <0.01, two-side t-test compared with the biolysis method; all experiments were performed in triplicate; error bar, mean ± SD.

### Precise comparison of released amino acids by ultra-high performance liquid chromatography-tandem mass spectrometry (UHPLC-MS/MS)

It is noteworthy that the formol titration method utilized above is rough to quantify released amino-acids, due to the coexistence of phosphates, polypeptides, fatty acids, and carbohydrates, whose reactions are also acid or alkaline to eventually affect the titration endpoint (31). To reliably measure the contents of released amino acids, UHPLC-MS/MS, which was demonstrated to be rapid, specific, and sensitive for profiling free amino acids (32, 33), was accordingly applied. All 20 amino acids except cysteine were detected in all SP products (Fig. 4A). Glutamic acid, the highest-abundance (∼8.386%) amino acid in SP (14), was highly abundant (951.03 ± 109.54 ug/ml) in the SP products of all 4 methods without statistical differences (Fig. 4A), while the second most abundant (∼5.793%) aspartic acid in SP (14), the other acidic amino acid, was liberated in relatively lower amounts, suggesting that the aforementioned pH falls (Fig. S2) were possibly due to the liberated glutamic acid mainly and aspartic acid partially. Nonetheless, the biolysis method released about 2.80, 3.01, and 3.64 times the amount of aspartic acid liberated by alternate freezing and thawing, water bath, and sonication respectively. Serine, the amino acid reported to be consumed first (34), was insignificantly liberated by the 4 methods. The releases of all remaining amino acids except tyrosine and arginine were significantly higher in our biolysis method than in other methods (Fig. 4A). However, arginine was reported to be a poor nitrogen source (9), and the medium without tyrosine alone or in combination with cysteine was reported to achieve denser culture than medium with all 20 amino acids (35), accounting for the growth conspicuousness of SP broth generated by the biolysis method than those by the other 3 methods (Fig. 2B and 2E). Collectively, the level of total amino acids released by the biolysis method was significantly higher in comparison to other methods (Fig. 4B), in line with the findings of formol titration assays (Fig. 2A), supporting that the biolysis method degrades SP more effectively than the 3 other methods.

**Fig. 4.**
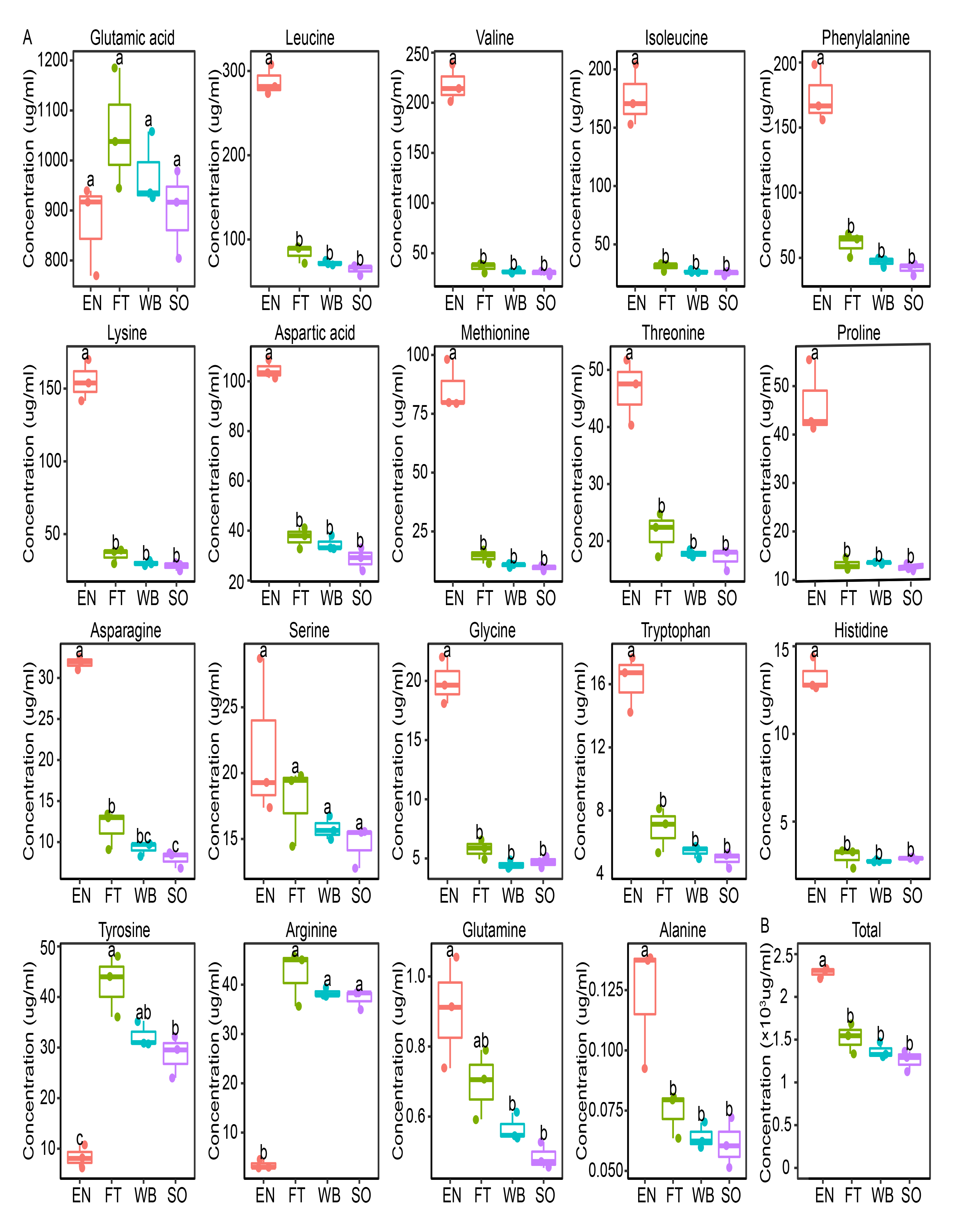
UHPLC-MS/MS profile of individual or total amino acids liberated by the 4 methods. A, for 19 single amino acids; B, for total amino acids. Different low-case letters indicate statistical difference at *P* <0.05, one-way ANOVA followed by Tukey’s post hoc test (n =3 in each group); EN, the biolysis method; FT, the alternate freezing and thawing; WB, the water bath method; SO, the sonication method. The total amount of free amino acids was calculated by summing concentrations of all detected amino acids.

### Further improved growth performance by formulating OSP into MSP

To further increase growth performance, a systematic investigation on OSP modification was conducted and finally, the formulated MSP was obtained to contain 15% (w/v) OSP, 2.524 g of KH_2_PO_4_, and ∼3.737 g of K_2_HPO_4_ (potassium phosphate buffer at pH 7.0, PPB7.0, totaling 0.04 M), 1% (w/v) glucose, and 2.03 mM MgSO_4_. It is noteworthy that the OSP became gradually alkaline over the course of the cultivation (Fig. 5A), possibly due to the excretion of the ammonia derived from the consumption of amino acids in OSP (36). On the contrary, an initial drop of the pH down to 6.6 was observed in MSP (Fig. 5A), possibly due to acid generation by glucose metabolism (37), as glucose was preferentially catabolized *via* carbon catabolic repression (38); then the pH increases gradually up to a maximum of 7.2 at 120 h (Fig. 5A), possibly due to the buffering capacity of PPB7.0 against alkalization generated by amino acid metabolism (Fig. S4A) (36). Together, MSP maintained pH in the best range of 6.5-7.5 as mentioned previously for *E. coli* cultivation (39), which is very important for microbial growth.

**Fig. 5.**
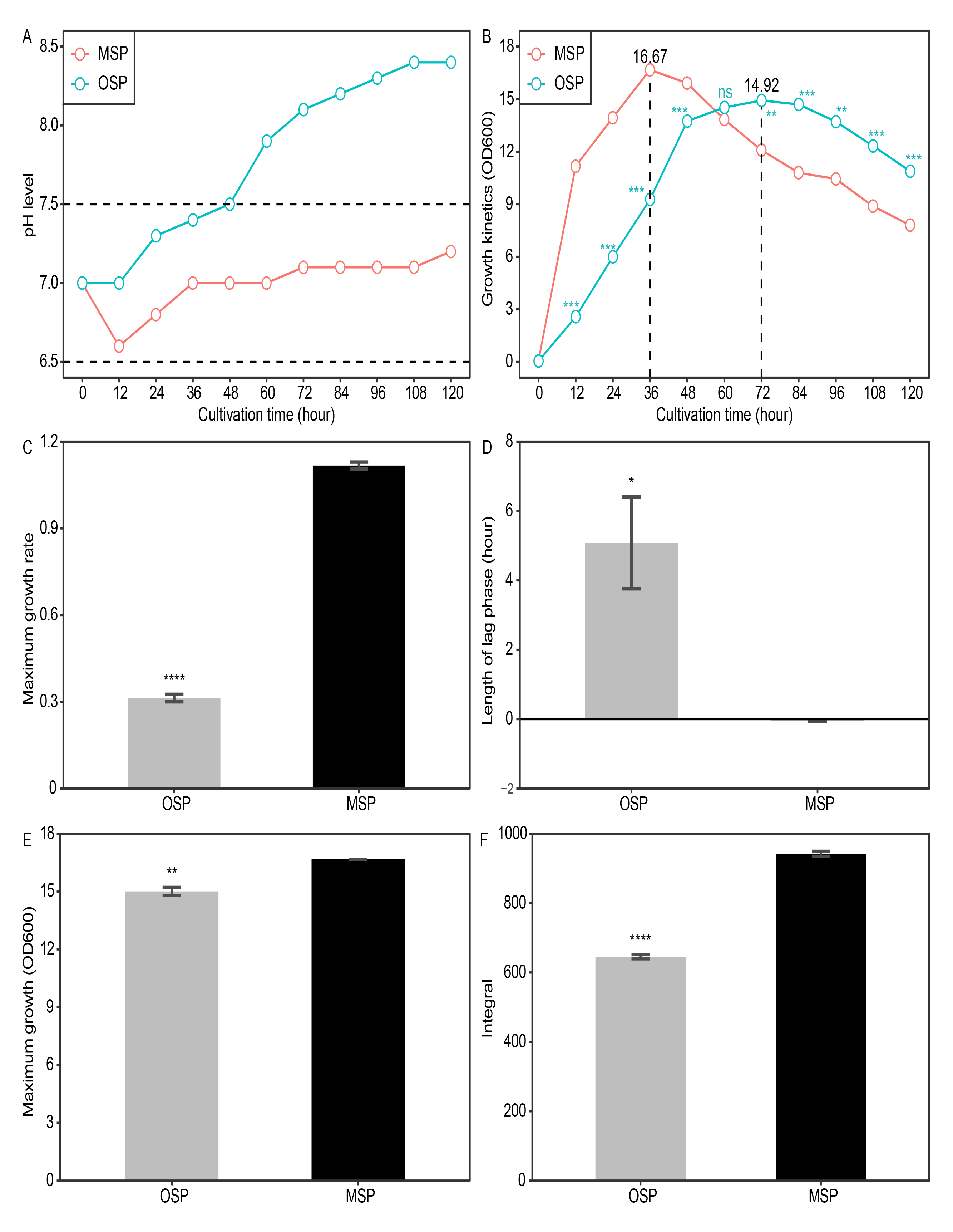
Improved growth performance by formulating OSP into MSP. pH maintenance (A) and growth curves (B), with 4 growth parameters including maximum growth rate (C), length of lag phase (D), maximum growth (E), and integral (F). Values and error bars represent means ± standard deviations from biological triplicate cultivations. \**P* <0.05, \*\**P* <0.01, \*\*\**P* <0.001, \*\*\*\**P* <0.0001, and ns, *P* >0.05, t-test compared with MSP. The best pH range of 6.5-7.5 for *E. coli* cultivation (39), as indicated by the dashed horizontal lines in panel A. The integral parameter shown here was based on the cultivation until 72 h, as indicated by the dashed vertical line in panel B.

Growth comparison showed that MSP increases the maximum OD600 from ∼14.92 to ∼16.67 (Fig. 5B), with a significant difference demonstrated by *grofit* extraction (Fig. 5E), possibly due to the addition of both PPB7.0 partially (Fig. S4B) and 2.03 mM MgSO_4_ mainly (Fig. S6A and Fig. S6D). This is because MgSO_4_ supplementation in the millimolar range harbors promoting effect on growth (35), and PPB7.0 maintained pH in the best range of 6.5-7.5 (39). Interestingly, the time to reach the maximum OD600 was reduced from 72 h to 36 h, attributing to a significantly increased maximum growth rate (Fig. 5C) in combination with a statistically reduced lag time (Fig. 5D). The increased maximum growth rate was due to the supplementation of PPB7.0 (Fig. S4C), 1% glucose (Fig. S5B), and 2.03 mM MgSO_4_ (Fig. S6B), and the reduced lag time was due to the amendment of 1% glucose (Fig. S5C), while PPB7.0 even increased the lag time (Fig. S4D). Glucose is one of the most favorable carbon sources to support the fast growth of *E. coli* when ammonia rather than a single poor nitrogen source such as arginine, glutamate, or proline is used as a nitrogen source (9). Our OSP has plentiful amino acids (Fig. 4) to generate sufficient ammonia, while glucose was negligible in all SP products of the 4 degradation methods due to below the analytical detection limit of UHPLC-MS/MS (data not shown). Therefore, glucose fuels growth and reduces lag time. However, the mechanism underlying why PPB7.0 increased both the maximum growth rate and the lag time awaits further investigation. Therefore, the integral value of MSP was significantly larger than that of OSP for the growth curves until 72-h cultivation (Fig. 5F), with the aim of achieving HCDC here.

### Outstanding growth performance in comparison to LB

For comparison, *E. coli* K-12 was cultivated in the reference medium LB. We found that MSP achieved a high averaged maximum growth (OD600) of ∼16.67, whilst only ∼ 6.30 for LB (Fig. 6A). Besides, MSP required only 36 h to reach maximum growth, much faster than LB which required 48 h (Fig. 6A), which is greatly beneficial for both basic research and industrial application. Extraction of 4 growth parameters showed that MSP achieved a significantly higher maximum growth rate (Fig. 6B), initiated growth immediately after inoculation (indicated as lag time <0 h) albeit with a significantly longer lag phase (Fig. 6C), and reached a higher maximum growth of 1.12 ± 0.01 h^−1^ (Fig. 6D) and integral (Fig. 6E), compared with LB. Collectively, MSP is superior to LB, attaining HCDC in batch culture.

**Fig. 6.**
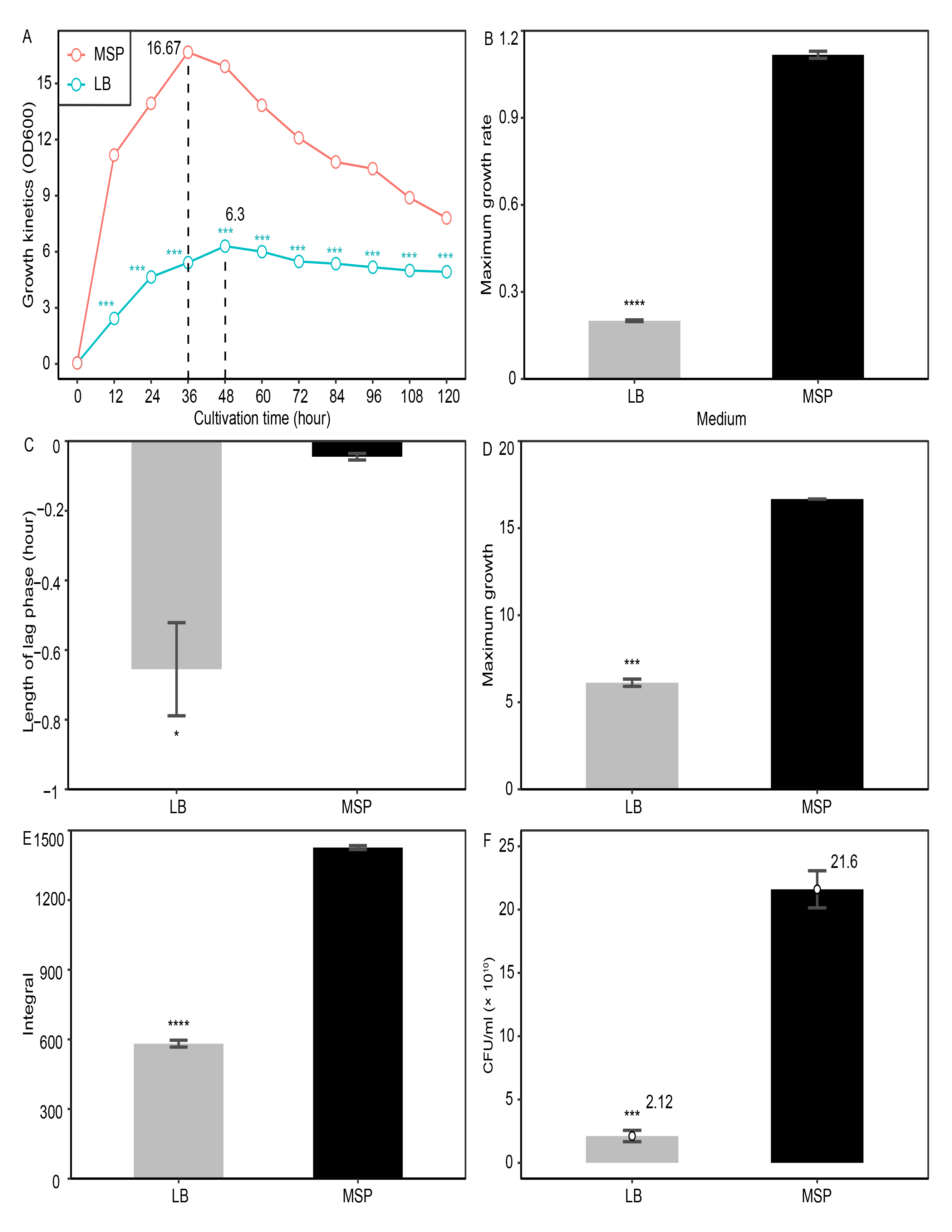
Outstanding growth performance of MSP in comparison to LB. Growth curves (A), maximum growth rate (B), length of lag phase (C), maximum growth (D), integral (E), and the amounts of viable cells cultivated at 36 h. Values and error bars represent means ± standard deviations from biological triplicate cultivations. \**P* <0.05, \*\*\**P* <0.001 and \*\*\*\**P* <0.0001, t-test compared with MSP. Dashed vertical lines in panel A indicate the times to reach the maximum OD600.

It was reported that cells in HCDC contain viable cells, viable but not culturable cells, cells under lysis, and dead cells (40). Among them, viable cells are more valuable. Therefore, viable cells were counted by plating cultures sampled at 36 h. Remarkably, MSP attained 2.16 × 10^11^ ml^-1^ of viable cells, ∼10.19 times the amount obtained by LB (Fig. 6F), while their OD600 fold is only ∼3.08 (16.67/5.42) (Fig. 6A). This indicated that MSP is more suitable to produce/maintain viable cells than LB even in the exponential growth phase. In conclusion, MSP attains HCDC with high viability.

## Discussion

Many effects have been made to search for new sources for preparing media (20, 41-43). However, none of them achieved HCDC. SP, which was demonstrated as a novel material for media preparation (20), was considered by us here as an ideal material to potentially develop culture media for HCDC (see detailed reasons, please refer to the “Introduction” section); this hypothesis was demonstrated in this study. To the best of our knowledge, it is the first time for us to demonstrate that SP can be utilized to achieve HCDC in batch mode for *E. coli* (20). Besides, so far, SP is the first material that can be developed to attain batch HCDC for *E. coli*.

To reach HCDC, investigations had been shifted from batch culture to several promising culture technologies, including fed-batch culture (2, 3), dialysis culture (4-6), and semicontinuous culture (7). However, these systems were not preferable in laboratories and even in industries, due to the requirement of sophisticated instrumentation or great waste of nutrients (dialysis culture as an example). In contrast, this study showed that medium development and modification can achieve HCDC in batch mode, highlighting the importance of the rebirth of medium development and modification.

OSP has plentiful free amino acids (Fig. 4) and short peptides (Fig. 3). Amino acids including both free and peptide-bound amino acids can be used as single carbon sources (44) and key substrates for achieving a high specific growth rate for *E. coli* (45), while their metabolisms result in considerably less acetate acid (46), which is detrimental for growth (11). Also, it is noteworthy that OSP contains negligible glucose, due to below the analytical detection limit of UHPLC-MS/MS (data not shown), avoiding the generation of acetate acid. The excellent growth performance (∼14.92 OD600) of OSP obtained in this study indicates that metabolism optimization *via* supplying “good” media can reach HCDC in batch mode, further supporting the rebirth of medium development and modification for HCDC in batch mode and the shift from developing new culture technologies to medium development and modification for HCDC in batch mode.

Finally, it is the first time for us to report a biolysis method with high efficiency for SP here (20). Based on its SP hydrolysates, MSP was formulated in this study and attained HCDC for *E. coli*. However, we realize that OSP or MSP with rich nutrients developed here could be extended to other applications, including recombinant protein production, DNA transformation, rehydration of lyophilized cells (20) and rapid resuscitation of lactic acid-induced sublethal injuries (47). Consequently, the biolysis method as well as OSP or MSP are fundamental, due to their great potential for wide application. Further studies should focus on exploring their extended applications.

## Materials and Methods

### Growth condition and growth kinetics for *B. subtilis*

Each aliquot of 1 ml *B. subtilis* fresh pre-cultures (∼1.00 OD600) in Nutrient Broth (NB) was inoculated into 10 ml fresh NB or LB and cultivated for 72 h (37℃, 200 rpm). To establish growth kinetics, OD600 measurement was periodically recorded by a UV–2700 spectrophotometer (Shimadzu, Japan) in triplicate at 12-h intervals. After 72- h cultivation, the supernatants of *B. subtilis* in ∼9-ml quantities were collected *via* centrifugation at 10,000 g for 20 min and preserved at 4℃ for downstream SP degradation.

### SP degradation by cultivation supernatant of *B. subtilis*

SP dry powder of 45 g was soaked in 300 ml distilled water and then autoclaved at 121℃ for 15 min to eliminate the effect of microorganisms on SP consumption. Our checking found that the initial SP solutions were at pH ∼7. After cooling down, 1 ml, 3 ml, and 9 ml of preserved NB or LB cultivation suspension of *B. subtilis* were added. At 5-h intervals, samples were collected for downstream formol titration and their pH levels were electrometrically monitored offline with a pH electrode (PHS-3C, INESA Scientific Instrument Co., Ltd, Shanghai, China). To eliminate the effect of different pH falls on SP degradation, all resulting SP solutions were adjusted to pH 7.0 with 1 M NaOH. Then, 1.893 g of KH_2_PO_4_ and 2.803 g of K_2_HPO_4_ (totaling 0.1 M potassium phosphate, pH 7.0) were jointly added to provide buffering capacity, followed by the addition of fresh cultivation supernatant of *B. subtilis* with the same volumes. After 4 successive addition-degradation cycles, the final products were centrifuged at 10,000 g for 20 min to obtain their aqueous supernatants. Then, to compensate for the volume loss, an appropriate amount of water was added to respective samples, which were subsequently neutralized by 1 M NaOH and finally sterilized by autoclaving at 121℃ for 15 min and preserved at 4℃ for the following experiments.

### Degradation procedures of the mechanical methods

Dried SP power was soaked in distilled water at 15% (w/v), autoclaved at 121℃ for 15 min, and subjected to processing by the following reference methods. For the sonication method, the SP solution was sonicated with a sonotrode of 2 mm diameter and 100% power input in 8 × 5 min periods without any cooling breaks in between. For the water bath method, the SP solutions were preheated at 60℃ for 30 min to avoid the break of the flask followed by a bath at 100℃ for 60 min. For the alternate freezing and thawing method, the SP liquids were processed by four freezing (-20℃) and thawing (room temperature) cycles. After processing, all samples were centrifuged at 10,000 g for 20 min, precipitates were discarded and the aqueous supernatants were sterilized by autoclaving at 121℃ for 15 min for the following experiments.

### Rough measurement of SP protein hydrolysis by formol titration

An Aliquot of 2 ml above sampled SP degradation solution after centrifugation was measured out with a pipette into each of two 100-ml Erlenmeyer flasks. To each Erlenmeyer flask added 5 ml of distilled water. One of the Erlenmeyer flasks served as a color screen and the other one severed as the tested sample. Besides, a third Erlenmeyer flask containing only 7 ml distilled water was used as a formalin blank. Five drops of the color indicator Phenolphthalein and then 2 ml of formalin were added into each of the three Erlenmeyer flasks. After amply twirling, a burette of 0.1 M NaOH was used for titration until the pink color of the indicator was visible. The titration of a sample subtracted by the titration of the formalin blank was then used to indicate the protein hydrolysis of this sample. In total, three biological replicates were conducted for each sampled SP degradation solution.

### Gel electrophoresis and densitometry by ImageJ

The polyacrylamide gel was composed of a stacking gel (30% acrylamide in 1.00 M Tris-HCl buffer, pH 6.8) and a 15% acrylamide separating gel in 1.5 M Tris-HCl buffer, pH 8.8. The supernatants obtained above were used to compare the degradation efficiency of the 4 methods. SDS-PAGE was performed under reducing conditions at 60 V through the stacking gel and 80 V in the separation gel until the tracking dye migrated to the bottom of the gel. Gel was stained in a solution of 1% (w/v) Coomassie blue R- 250, 40% (v/v) methanol, and 16.6% (v/v) acetic acid, and decolorized under continuous shaking in a 25% (v/v) ethanol solution containing 10% (v/v) acetic acid. A low-molecular-weight pre-stained protein ladder (Affinity Biosciences Group Ltd., Jiangsu, China) composed of ten recombinant protein bands of 250, 150, 100, 70, 50, 40, 35, 25, 20, 15, and 10 kDa was used to indicate the molecular weight. Gels were scanned with AlphaImager HP Gel Imaging System (Alpha Innotech Corp., USA) under UV epi-illumination. Images were saved as a PNG file with a size of 16-bit and a resolution of 1,228 × 884 pixels and then converted to 8-bit grayscale images using the ImageJ (Sun Microsystems Inc., USA) for densitometric quantification. After background subtraction, the area intensities of <10 kDa, 10-15 kDa, 15-20 kDa, and >20 kDa were individually obtained to show degradation effectiveness.

### Quantification of released amino acids by UHPLC-MS/MS

#### Sample Preparation

Supernatant samples of approximately 8 ml of each obtained by the above 4 methods were sent in triplicate on dry ice to Shanghai Profleader Biotech Co., Ltd (Shanghai, China). Upon arrival, 100 µl of each sample was immediately added to 400 µl of 80% methanol (Shanghai Ampere Laboratory Technology Co., Ltd., Shanghai, China) and allowed to stand for 30 min at 40 °C, followed by centrifugation at 15,000 g and 4 °C for 15 min to precipitate the proteins. Afterward, the supernatant was isometrically mixed with the internal standards labeled with stable isotopes (Sigma-Aldrich, Shanghai, China) for UHPLC-MS/MS analysis.

#### UHPLC-MS/MS analysis

Simultaneous quantification of 20 free amino acids without any derivatization was conducted on an Agilent 1290 Infinity II UHPLC system coupled to a 6470A Triple Quadrupole mass spectrometry (Santa Clara, CA, United States). Samples were pumped into an ACQUITY UPLC® BEH Amide column from Waters (Milford, USA) with a dimension of 100 × 2.1 mm, a particle size of 1.7 μm and a porosity of 130 Å at a flowrate of 0.2 mL/min. The mobile solution (A) consisted of 10 mM ammonium acetate (LC-MS grade) and (B) 90% acetonitrile. The chromatographic separation was carried out with a flowrate of 0.2 ml/min, a column temperature of 40°C, and a gradient elution program as follows: 0-1 min = 90%, 4 min = 85%, 12 min = 70%, 24-16 min = 50%, and 16.1-20 min = 90%.

The eluted analysts were then directly ionized with an electrospray ionization source in positive mode with the following operating parameters: drying gas flow (5 L/min), drying gas temperature 300°C, nebulizer gas (45 psi), sheath gas flow (11 L/min), sheath gas temperature (350°C), and capillary voltage (4000 V). The detection was achieved with an Agilent electron multiplier in dynamic multiple reaction monitoring (dMRM) mode. The optimized dMRM transition (precursor -> product), fragmentor voltage, and collision energy (CE) were shown in Table S1. The Agilent MassHunter software (version B.08.00) was applied for data acquisition, while the MassHunter Workstation Software (version B.08.00, default parameters) was employed for data processing under manual inspection to guarantee accurate measurements. External calibration in the range of 0.01-5 μg/mL *via* serially diluting a working solution of the standards of 20 amino acids was utilized for quantification.

### Batch cultivation and growth measurement

About 5 ml of *E. coli* fresh pre-culture was inoculated into 100 ml of the appropriate medium in a 500-ml Erlenmeyer flask, and incubated at 37°C with orbital agitation of 200 rpm, the highest speed that produced negligible foaming. For monitoring cell growth, successive samples from shake flask cultures at 12-h intervals from 12 h to 120 h were collected to measure OD600 in cuvettes with a UV–2700 spectrophotometer (Shimadzu, Japan). pH was monitored offline with a pH electrode (PHS-3C, INESA Scientific Instrument Co., Ltd, Shanghai, China) if required. When measuring OD600, 3 ml of each culture was centrifuged for 2 min at 10,000 g, the supernatant was removed, and the pellet was resuspended in sterile phosphate-buffered saline (PBS), followed by serial dilutions with PBS when appropriate to remain within the range of linearity of the instrument for OD600 measurement.

### Extracting 4 growth characteristics from growth data

The *grofit* R package (version 1.1.1) (48) was employed to in-depth characterize the growth curves. First, model-free smoothing spline was selected for fitting the resultant growth data of each replicate, considering that spline estimates growth parameters more accurately than model-based fits (48). Then, the fitted growth curve was utilized to estimate 4 typical growth parameters as follows: the length of lag phase (hour), the maximum instantaneous growth rate represented by the maximum slope, the maximum growth (OD600), and the integral (area under the curve) estimated by numerical integration (49).

### MSP preparation and growth evaluation

Around 2.524 g of KH_2_PO_4_, ∼3.737 g of K_2_HPO_4_, 10 g of glucose, and 50 mg (∼ 2.03 mM) of MgSO_4_.7H_2_O were added into 1 L of centrifuged OSP. The resulting broth MSP was then filter-sterilized by using a 0.22-μm membrane filter unit (Millex®-GP, Merck Millipore Ltd. Tullagreen, Carrigtwohill, Co. Cork, Ireland). Growth experiments and extraction of 4 growth parameters were conducted to show which buffered OSP achieved the best performance as described above.

### Reference cultivation in LB medium

For comparison, *E. coli* K-12 was cultivated in LB medium based on the protocol as described above. Then, 4 growth parameters were extracted from the growth data of each replicate by using *grofit* (version 1.1.1) (48) according to the protocol described above.

### Viable cell enumeration

Aliquots (1 ml) of the samples collected from MSP or LB cultivation at 36 h were serially diluted 10-fold by using 9 ml of each corresponding broth. Then, 0.1 ml of adequately diluted aliquots were aseptically pipetted onto the center of LB plates in triplicate and spread uniformly by a sterile glass rod. All inoculated plates were incubated at 37°C for 24 h and the number of colony-forming units was visually enumerated. The plated dilutions were selected so that about 30-300 viable cells were spread on each plate.

### Statistical analysis

Data were presented as means ± SD (standard deviations) of biological triplicate experiments. Differences between multiple groups were analyzed by one-way analysis of variance (ANOVA) followed by Tukey’s post hoc test with a *P* value of <0.05 considered to be statistically significant, while pairwise differences were performed by the Student’s t-test. All statistical analyses were performed by using R.

## Acknowledgments

This work was jointly supported by the Natural Science Foundation of Guangxi ([grant number 2020GXNSFAA159032]; and the Natural Science Foundation of China [grant number 32060144] for Yizhuang Zhou.

## References

1. Shiloach J, Fass R. 2005. Growing E. coli to high cell density--a historical perspective on method development. Biotechnol Adv 23:345–57.

2. Teworte S, Malcı K, Walls LE, Halim M, Rios-Solis L. 2022. Recent advances in fed-batch microscale bioreactor design. Biotechnol Adv 55:107888.

3. Ramírez N, Ubilla C, Campos J, Valencia F, Aburto C, Vera C, Illanes A, Guerrero C. 2021. Enzymatic production of lactulose by fed-batch and repeated fed-batch reactor. Bioresour Technol 341:125769.

4. Märkl H, Zenneck C, Dubach AC, Ogbonna JC. 1993. Cultivation of Escherichia coli to high cell densities in a dialysis reactor. Appl Microbiol Biotechnol 39:48–52.

5. Pörtner R, Märkl H. 1998. Dialysis cultures. Appl Microbiol Biotechnol 50:403–14.

6. Fuchs C, Köster D, Wiebusch S, Mahr K, Eisbrenner G, Märkl H. 2002. Scale-up of dialysis fermentation for high cell density cultivation of Escherichia coli. J Biotechnol 93:243–51.

7. Quinlan AV. 1986. A semicontinuous culture model that links cell growth to extracellular nutrient concentration. Biotechnol Bioeng 28:1455–61.

8. Sezonov G, Joseleau-Petit D, D’Ari R. 2007. Escherichia coli physiology in Luria-Bertani broth. J Bacteriol 189:8746–9.

9. Bren A, Park JO, Towbin BD, Dekel E, Rabinowitz JD, Alon U. 2016. Glucose becomes one of the worst carbon sources for E.coli on poor nitrogen sources due to suboptimal levels of cAMP. Sci Rep 6:24834.

10. Riesenberg D. 1991. High-cell-density cultivation of Escherichia coli. Current Opinion in Biotechnology 2:380–384.

11. Pinhal S, Ropers D, Geiselmann J, de Jong H. 2019. Acetate Metabolism and the Inhibition of Bacterial Growth by Acetate. J Bacteriol 201.

12. Lupatini AL, Colla LM, Canan C, Colla E. 2017. Potential application of microalga Spirulina platensis as a protein source. J Sci Food Agric 97:724–732.

13. Ishimi Y, Sugiyama F, Ezaki J, Fujioka M, Wu J. 2006. Effects of spirulina, a blue-green alga, on bone metabolism in ovariectomized rats and hindlimb-unloaded mice. Biosci Biotechnol Biochem 70:363–8.

14. Seyidoglu N, Inan S, Aydin C. 2017. A Prominent Superfood: Spirulina platensis. Superfood and Functional Food - The Development of Superfoods and Their Roles as Medicine doi:DOI: 10.5772/66118.

15. Chaiklahan R, Chirasuwan N, Triratana P, Loha V, Tia S, Bunnag B. 2013. Polysaccharide extraction from Spirulina sp. and its antioxidant capacity. Int J Biol Macromol 58:73–8.

16. Li TT, Huang ZR, Jia RB, Lv XC, Zhao C, Liu B. 2021. Spirulina platensis polysaccharides attenuate lipid and carbohydrate metabolism disorder in high-sucrose and high-fat diet-fed rats in association with intestinal microbiota. Food Res Int 147:110530.

17. Doelle HW, Ewings KN, Hollywood NW. Regulation of glucose metabolism in bacterial systems, p 1–35. *In* (ed), Springer Berlin Heidelberg,

18. de Caire GZ, Parada JL, Zaccaro MC, de Cano MMS. 2000. Effect of Spirulina platensis biomass on the growth of lactic acid bacteria in milk. World Journal of Microbiology and Biotechnology 16:563–565.

19. Wu LC, Ho JA, Shieh MC, Lu IW. 2005. Antioxidant and antiproliferative activities of Spirulina and Chlorella water extracts. J Agric Food Chem 53:4207–12.

20. Kheirabadi E, Macia J. 2022. Development and evaluation of culture media based on extracts of the cyanobacterium Arthrospira platensis. Front Microbiol 13:972200.

21. Jeong Y, Choi WY, Park A, Lee YJ, Lee Y, Park GH, Lee SJ, Lee WK, Ryu YK, Kang DH. 2021. Marine cyanobacterium Spirulina maxima as an alternate to the animal cell culture medium supplement. Sci Rep 11:4906.

22. Stewart DE, Farmer FH. 1984. Extraction, identification, and quantitation of phycobiliprotein pigments from phototrophic plankton. Limnology and Oceanography 29:392–397.

23. Lin R, Liu W, Piao M, Zhu H. 2017. A review of the relationship between the gut microbiota and amino acid metabolism. Amino Acids 49:2083–2090.

24. Zhao L, Ye B, Zhang Q, Cheng D, Zhou C, Cheng S, Yan X. 2019. Construction of second generation protease-deficient hosts of Bacillus subtilis for secretion of foreign proteins. Biotechnol Bioeng 116:2052–2060.

25. Margot P, Karamata D. 1996. The wprA gene of Bacillus subtilis 168, expressed during exponential growth, encodes a cell-wall-associated protease. Microbiology (Reading) 142 (Pt 12):3437–44.

26. Rutherfurd SM. 2010. Methodology for determining degree of hydrolysis of proteins in Hydrolysates: a review. J AOAC Int 93:1515–22.

27. Pez Jaeschke D, Rocha Teixeira I, Damasceno Ferreira Marczak L, Domeneghini Mercali G. 2021. Phycocyanin from Spirulina: A review of extraction methods and stability. Food Res Int 143:110314.

28. Tavanandi HA, Mittal R, Chandrasekhar J, Raghavarao KSMS. 2018. Simple and efficient method for extraction of C-Phycocyanin from dry biomass of Arthospira platensis. Algal Research 31:239–251.

29. Furuki T, Maeda S, Imajo S, Hiroi T, Amaya T, Hirokawa T, Ito K, Nozawa H. 2003. Rapid and selective extraction of phycocyanin from Spirulina platensis with ultrasonic cell disruption. Journal of Applied Phycology 15:319–324.

30. Jing Y, Mu C, Wang H, Shen J, Zoetendal EG, Zhu W. 2022. Amino acid utilization allows intestinal dominance of Lactobacillus amylovorus. Isme j 16:2491–2502.

31. Brown JH. 1923. The Formol Titration of Bacteriological Media. J Bacteriol 8:245–67.

32. Jin Y, Yang Wang C, Hu W, Huang Y, Li Xu M, Wang H, Kong X, Chen Y, Dong TT, Qin Q, Keung Tsim KW. 2019. An optimization of ultra-sonication-assisted extraction from flowers of Apocynum venetum in targeting to amount of free amino acids determined by UPLC-MS/MS. Food Quality and Safety 3:52–60.

33. Weber P. 2022. Determination of amino acids in food and feed by microwave hydrolysis and UHPLC-MS/MS. J Chromatogr B Analyt Technol Biomed Life Sci 1209:123429.

34. Yang Y, A MP, Höfler C, Poschet G, Wirtz M, Hell R, Sourjik V. 2015. Relation between chemotaxis and consumption of amino acids in bacteria. Mol Microbiol 96:1272–82.

35. Studier FW. 2005. Protein production by auto-induction in high density shaking cultures. Protein Expr Purif 41:207–34.

36. Krause M, Ukkonen K, Haataja T, Ruottinen M, Glumoff T, Neubauer A, Neubauer P, Vasala A. 2010. A novel fed-batch based cultivation method provides high cell-density and improves yield of soluble recombinant proteins in shaken cultures. Microb Cell Fact 9:11.

37. Rosano GL, Ceccarelli EA. 2014. Recombinant protein expression in Escherichia coli: advances and challenges. Front Microbiol 5:172.

38. Magasanik B. 1961. Catabolite repression. Cold Spring Harb Symp Quant Biol 26:249–56.

39. Scheidle M, Dittrich B, Klinger J, Ikeda H, Klee D, Büchs J. 2011. Controlling pH in shake flasks using polymer-based controlled-release discs with pre-determined release kinetics. BMC Biotechnol 11:25.

40. Andersson L, Strandberg L, Enfors SO. 1996. Cell segregation and lysis have profound effects on the growth of Escherichia coli in high cell density fed batch cultures. Biotechnol Prog 12:190–5.

41. Hoover SW, Marner WD, 2nd, Brownson AK, Lennen RM, Wittkopp TM, Yoshitani J, Zulkifly S, Graham LE, Chaston SD, McMahon KD, Pfleger BF. 2011. Bacterial production of free fatty acids from freshwater macroalgal cellulose. Appl Microbiol Biotechnol 91:435-46.

42. Batista KA, Fernandes KF. 2015. Development and optimization of a new culture media using extruded bean as nitrogen source. MethodsX 2:154–8.

43. Narh C, Frimpong C, Mensah A, Wei Q. 2018. Rice Bran, an Alternative Nitrogen Source for Acetobacter xylinum Bacterial Cellulose Synthesis. BioResources 13.

44. Burkovski A, Krämer R. 2002. Bacterial amino acid transport proteins: occurrence, functions, and significance for biotechnological applications. Appl Microbiol Biotechnol 58:265–74.

45. Maser A, Peebo K, Vilu R, Nahku R. 2020. Amino acids are key substrates to Escherichia coli BW25113 for achieving high specific growth rate. Res Microbiol 171:185–193.

46. Krautkramer KA, Fan J, Backhed F. 2021. Gut microbial metabolites as multi-kingdom intermediates. Nat Rev Microbiol 19:77–94.

47. Zeng M, Yang S, Meng L, Jia S, Zhou L, Lao X, Tan S, Zhou Y. 2023. Developing a de novo designed broth to rapidly recover lactic acid-injured Escherichia coli to ensure almost no multiplication during repair for precise enumeration. Food Control 153.

48. Kahm M, Hasenbrink G, Lichtenberg-Fraté H, Ludwig J, Kschischo M. 2010. grofit: Fitting Biological Growth Curves with R. Journal of Statistical Software 33:1–21.

49. Hasenbrink G, Kolacna L, Ludwig J, Sychrova H, Kschischo M, Lichtenberg-Fraté H. 2007. Ring test assessment of the mKir2.1 growth based assay in Saccharomyces cerevisiae using parametric models and model-free fits. Appl Microbiol Biotechnol 73:1212–21.

